# Predicting the temperature-driven development of stage-structured insect populations with a Bayesian hierarchical model

**DOI:** 10.1101/2022.10.04.510664

**Authors:** Kala Studens, Ben Bolker, Jean-Noel Candau

## Abstract

The management of forest pests relies on an accurate understanding of the species’ phenology. Thermal performance curves (TPCs) have traditionally been used to model insect phenology; many such models have been proposed and fitted to data from both wild and laboratory-reared populations, most of which have used maximum likelihood estimation (MLE). Analyses typically present point estimates of parameters with confidence intervals, but estimates of the correlations among TPC parameters are rarely provided. Neglecting aspects of model uncertainty such as correlation among parameters may lead to incorrect confidence intervals of predictions. This paper implements a Bayesian hierarchical model of insect phenology incorporating individual variation, quadratic variation in development rates across insects’ larval stages, and non-parametric adjustment terms that allow for deviations from a parametric TPC. We use Hamiltonian Monte Carlo (HMC) for estimation; the model is fitted to a laboratory-reared spruce budworm population as a case study. We assessed the accuracy of the model using stratified, 10-fold cross-validation. Using the posterior samples, we found prediction intervals for spruce budworm development for a given year.

## Introduction

Many efforts have been made to improve predictions of insect development and phenology in the past century (Uvarov, 1931). These studies were historically motivated by the management of insect pests (Crimmins et al., 2020; Pruess, 1983); more recently, climate change and the increasing threat from invasive species have renewed interest in incorporating quantitative models of insect development and phenology in process-based models of geographic distribution (Régnière et al., 2012), evolutionary ecology (Bewick et al., 2016), and biological invasion (Porter et al., 1991).

Insects accomplish their life cycle by developing through discrete morphological stages. Development rate depends on several climatic variables, of which temperature is considered the most important (Rebaudo and Rabhi, 2018). Most models of insect development rely on two principles: a *thermal performance curve* (TPC) that formalizes the relationship between development rate and temperature (Chuine and Régnière, 2017), and a *rate summation* that states that maturation to the next stage occurs when development (based on the TPC and the distribution of environmental temperature) reaches a threshold.

TPC models typically provide point estimates with standard errors or confidence intervals (CIs: Quinn, 2017; Rebaudo and Rabhi, 2018); estimates of parameter correlation are rarely given. While CIs are sufficient for inference on model parameters, downstream quantities such as development rate cannot be accurately assessed without knowing the correlation among parameter estimates. Development rates are often more salient than the parameters themselves, especially when TPCs are parameterized using models that are chosen for statistical convenience rather than biological interpretability. Most models of insect development are fitted using either non-linear least squares or maximum likelihood estimation (MLE: Damos and Savopoulou-Soultani, 2012). While one can use these methods to estimate correlations among parameters, and hence derive CIs for predictions (Bolker, 2008), Bayesian methods provide samples of the full multivariate posterior distribution and thus enable CIs for any function of model parameters. Bayesian methodology also allows the specification of informative prior distributions; we can use the extensive literature on insect phenology to develop appropriate priors (McCARTHY and Masters, 2005).Bayesian methods have been used to model insect herbivory (Lemoine et al., 2014) and plant phenology (Ceglar et al., 2011; Fu et al., 2012; Qiu et al., 2020; Thorsen and Höglind, 2010), but not for TPCs. Mc-Manis et al. (2018) impose Bayesian prior distributions on the parameters in their model, but the parameter estimation is done by minimizing the negative log posterior rather than sampling a full posterior distribution; this approach leverages prior knowledge, but does not provide the benefits of convenient error propagation.

In this study we developed a Bayesian hierarchical model of individual-based, temperature driven development for spruce budworm (*Choristoneura fumiferana*). The resulting posterior distribution was used to simulate one year of larval development, and to obtain prediction intervals for development achieved by a given date, or of dates by which a critical developmental threshold occurs. Using cross-validation, we compared the Bayesian fitting method to MLE, and compared a semi-parametric model to a strictly parametric model of insect development.

## Methods

### Study System

The eastern spruce budworm is native to the boreal forests of North America. During periodic outbreaks, its populations have caused extensive tree mortality, especially of white spruce and balsam fir trees (Blais, 1983). The species is univoltine; its six larval stages (*instars*) are delineated by the shedding of head capsules. Upon emergence from eggs in the summer, the first instar larvae disperse and form hibernacula (cocoon-like shelters) in which they moult to second instar and enter diapause, a resting stage in which they spend the duration of the winter. Second instar larvae emerge from these hibernacula in the early spring and feed on old needles until budburst. In the third to sixth larval instars, larvae feed on new foliage. The synchrony of these larval stages with new foliage is crucial for survival (Lawrence et al., 1997); understanding the timing of each phenomenon is necessary in order to predict the impact of climate change on the species’ behaviour. The pupal stage is typically reached in early summer, and moths emerge several days later. Understanding the timing of adult emergence is important for modelling the species’ landscape-scale dispersal (Sturtevant et al., 2013).

Early models of spruce budworm development neglected individual variability (Bean, 1961; Cameron et al., 1968; Dennis et al., 1986; Hudes and Shoemaker, 1988; Lysyk, 1989). Régnière (1984; 1987) was the first to account for variability in a two-step procedure that separately estimated the effect of temperature on median development rate and the distribution of individual rates around the median, modeling the rate using a Type I generalized logistic distribution (Balakrishnan and Leung, 1988). Stedinger et al. (1985) modeled the distribution of larvae in each larval stage as Dirichlet-multinomial distribution, accounting for environmental conditions at three spatial scales (regional, site and individual). More recently, Régnière et al. (2012) proposed a general framework for quantifying insect development that models individual variability as a log-Normal distribution.

### Data Collection

The data comes from a spruce budworm rearing experiment described in Wardlaw et al. (2022). Samples of diapausing larvae were collected as described in Candau et al. (2019) from Timmins, Ontario in accordance with the methods in Perrault et al. (2021). Upon emergence from diapause, individuals were collected and placed in separate containers containing artificial diet created to mimic the nutrition that the insects would consume in the wild. The development of each insect was observed daily; a moult was reported once a larva had shed its head capsule. The colony was divided into seven sub-populations, each of which was reared at a different constant temperature, at evenly spaced temperatures from 5 − 35^◦^C. Due to the potential for high mortality rates at temperatures outside a “sustainable” temperature range of 15 − 25^◦^C, an extra rearing step was taken for populations at the extreme temperatures. For each stage, the times to moult at the extreme temperatures were estimated using BioSIM (Régnière et al., 2014). The insects were held at the extreme temperatures for approximately half the predicted moulting time and then moved to 20^◦^C for the remainder of each stage. Each sub-population contained 250 individuals, resulting in a total of 1,750 individuals. Since the main objective of the project was to observe development rates in the larval instar stages, survival was not considered in the modelling process. Thus, the data used to fit the model consisted only of individuals who survived the full rearing process from the first larval instar to pupation (Figure 1). Only the first new generation from the wild population was used to fit the data, to eliminate any generational effect of laboratory rearing. The low survival rate in the 15^◦^C treatment is likely due to the fact that it is on the edge of the sustainable growth regime; the insects at this temperature may have benefited from the transfer treatment performed for the larvae exposed to more extreme temperatures.

**Figure 1:**
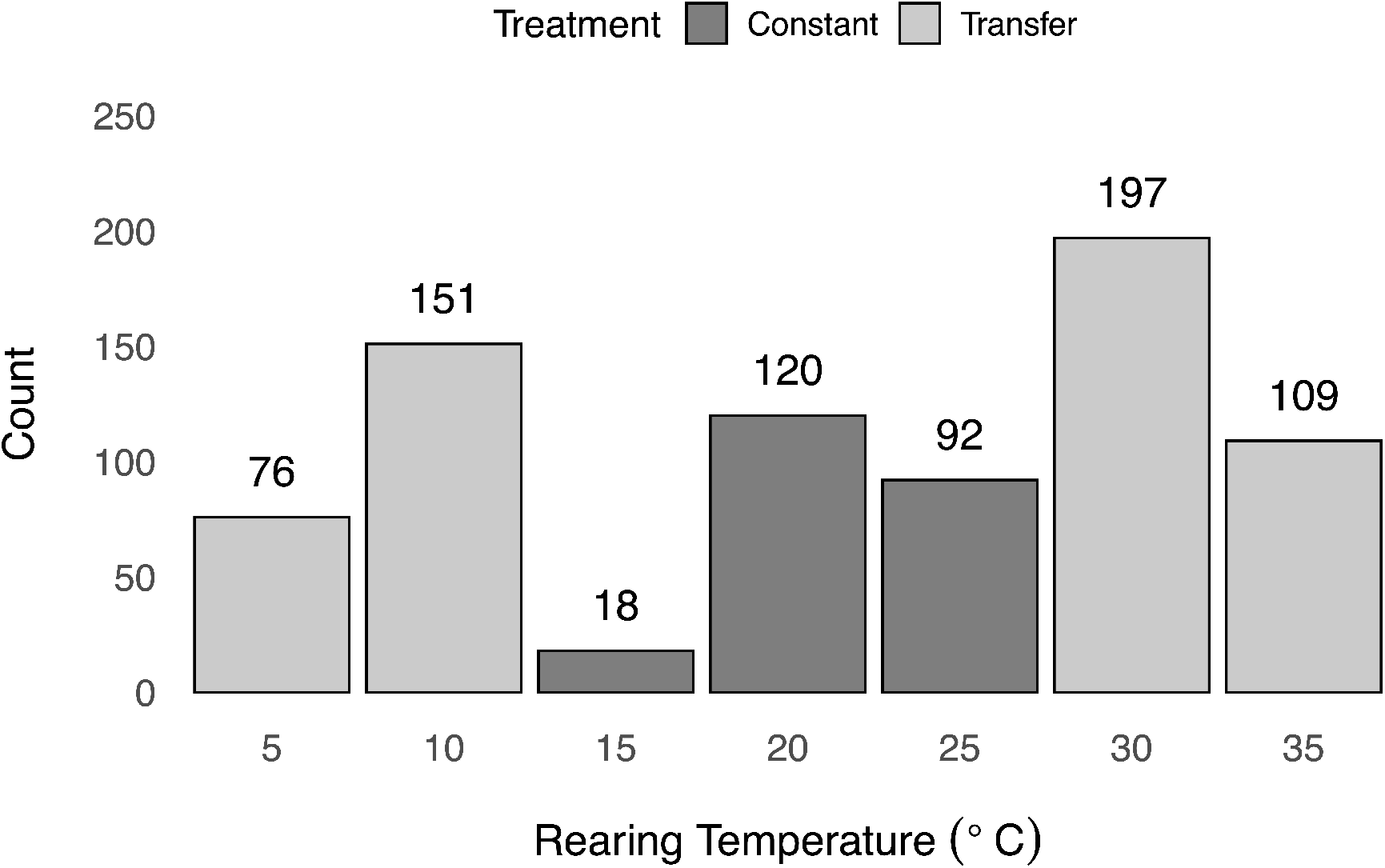
Surviving sub-population counts. Dark bars represent sustainable treatment temperatures, light bars represent extreme treatment temperatures.

### Likelihood Structure

In the model of Régnière et al. (2012), an individual’s physiological age *a* within a developmental stage is defined as the proportion of the stage that they have completed. The age of an individual *i* is represented as

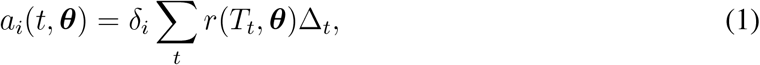

where *r*(*T*, ***θ***) is the population median development rate as a function of temperature *T* and the vector of development rate parameters ***θ***, and *δ*_*i*_ is a multiplier representing the development rate of individual *i*, relative to the population median; *δ*_*i*_ is independent of temperature. The multiplier Δ_*t*_ represents the time spent at temperature *T*_*t*_.

An age of 1.0 signals the end of a developmental stage. Therefore, if *t*_*m*_ is the amount of time taken by individual *i* to complete a stage, then *a*_*i*_(*t*_*m*_, ***θ***) = 1. Since observations were taken daily, we do not directly observe *t*_*m*_. Instead, the data are interval censored; we know only the one-day interval within which an individual completed each developmental stage (i.e. *t*_*m*_ *∈* [*t*_1_, *t*_2_]). Thus

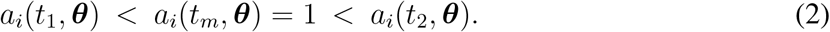

Substituting for Equation 1 and rearranging gives

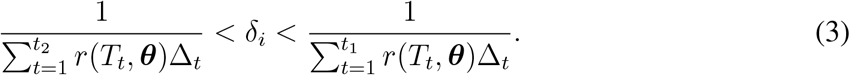

If we treat *δ*_*i*_ as a random variable with cumulative distribution *F*_*δ*_, then the likelihood of our observation of individual *i* given parameters ***θ*** is

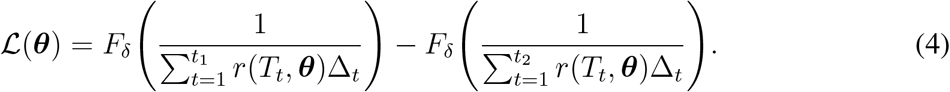

Régnière et al. (2012) assign a log-Normal distribution to *δ*_*i*_ to ensure that *δ >* 0 and that the distribution of individual variation is positively skewed. Their parameterization gives *E*(*δ*) = 1, thus centring the population mean at the development rate curve. Therefore, we have

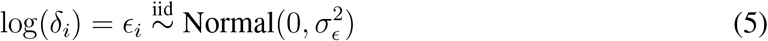

for each individual *i* = 1, …, *N*, where *σ*_*ϵ*_ is random. In this parameterization, we centre the distribution at the population median instead of the mean, so that a population’s development times and development rates have the same distribution across individuals (i.e.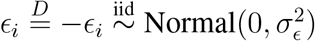).

Régnière et al. (2012) included a treatment-level, multiplicative random effect *υ*_*j*_, which allows the mean development times at the treatment temperatures to deviate from those predicted by the parametric model. This term adds flexibility to the model, relaxing the strict parametric form of the development rate curve to a semi-parametric form; the additional *non-parametric adjustment* parameters are included to decrease the systematic error caused by misspecification of the development rate curve. Régnière et al. (2012) specified 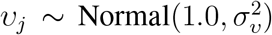 (where 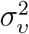 is a fitted parameter). In our case, because the magnitude of bias (degree of model misspecification) varied strongly with temperature, the Normal distribution did not allow enough flexibility, as indicated by the appearance of divergent transitions (Stan Development Team, 2020) and bimodality in the posterior distribution. To increase model flexibility, we changed the distribution of the random effects. (Changing the specification of a Bayesian model in order to overcome computational problems is widely practiced by modellers, and is explicitly recommended by leading Bayesian analysts (Gelman et al., 2020)). The *υ*_*j*_ parameters were redefined as

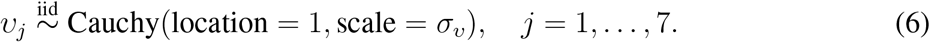

The resulting development rates are

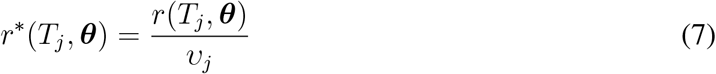

so the full log likelihood can be written as

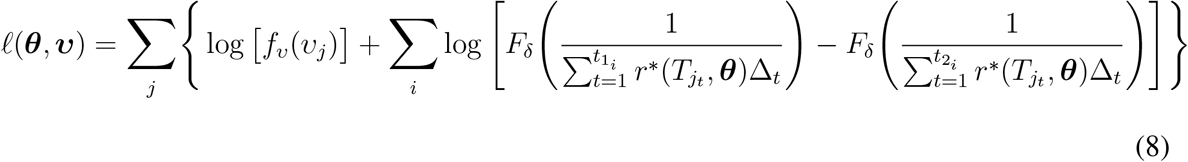

where *i* denotes individual, *j* denotes temperature treatments, and *f*_*υ*_ is the probability density function of a Cauchy distribution with location parameter 1 and scale parameter *σ*_*υ*_.

The observations for the L2 stage were interval censored, while the observations for the remaining stages were double censored (Calle, 2002). That is, the start time for the L2 stage is known but the exact end time is interval censored, while for the remaining stages both the start and end times are interval censored. Since observations were made daily, the interval for the true moult time *t* for an individual in the L2 stage is *t ∈* [*t*_obs_ − 1, *t*_obs_], while the interval for an individual in any other development stage is *t ∈* [*t*_obs_ − 1, *t*_obs_ + 1]. Here, *t*_obs_ is the observed moult time; for the L2 stage it is the number of days until a moult is observed, while for the remaining stages it is the number of days between the observations of successive moults.

### Development Rate Model

The Schoolfield et al. (1981) model with the reference temperature generalization presented in Ikemoto (2005) was chosen for the Bayesian implementation of the Régnière et al. (2012) likelihood framework. The development rate equation in this model, representing the rate of the enzyme-catalyzed reaction governing growth, is

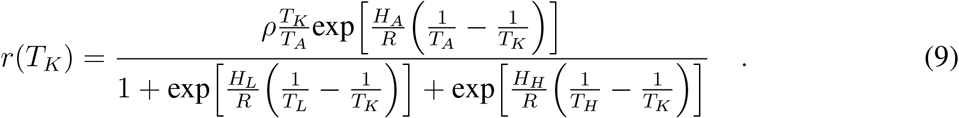

Here, *R* = 1.987 kcal · K^−1^ · mol^−1^ is the universal gas constant and *T*_*K*_ is the input temperature in degrees Kelvin, while the remaining values are the curve parameters; all temperatures in this model are represented in degrees Kelvin. The equation is based on the assumption that the enzymes controlling growth rate can be active; inactive due to low temperature; or inactive due to high temperature. These states are assumed to be reversible. The reciprocal of the denominator represents the fraction of enzymes in the active state. *T*_*A*_ is the temperature at which the fraction of active enzymes is maximized; the value of *ρ* in the numerator represents the development rate when *T*_*K*_ = *T*_*A*_. In the parameterization from Ikemoto (2005), the value of the reference temperature *T*_*A*_ is calculated from the other parameters as follows:

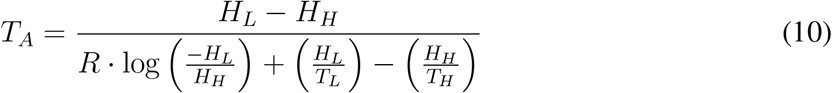

This development rate model provides some important advantages within a Bayesian framework. The first is the absence of temperature thresholds; while many of the development models in the literature include sharp temperature thresholds, there is little evidence that they are crucial to modelling development (Chuine and Régnière, 2017). While increased mortality does occur at extreme temperatures, mortality is a separate process from development. Under the current likelihood structure, individuals vary around a single population median development rate curve. Since the temperature thresholds would be features of this median curve, each individual would be assumed to have the same threshold. A fully continuous development rate curve that approaches but does not achieve zero at the extreme temperatures provides more flexibility, since development can effectively drop to zero for certain individuals while allowing others to continue development. McManis et al. (2018) deal with this behaviour by implementing additive rather than multiplicative variation around the mean development curve. However, this approach can result in negative development rate estimates, especially at temperatures for which development is already low.

The discontinuity of the rate curve in models when sharp temperature thresholds can cause computational problems, especially when using estimation methods that rely on following gradients of the log-likelihood surface. As well as making the surface non-differentiable in certain parameter regimes, these parameters also destabilize the log-likelihood. For instance, if a sampled lower temperature threshold is greater than a point where development occurs, the probability of the event becomes zero and the log-likelihood goes to negative infinity. This behaviour makes it difficult to specify priors for these threshold parameters.

### Stage Structure

The duration of the spruce budworm’s larval stages decreases from the second larval stage to the fourth and then increases until pupation (Régnière et al., 2012). Thus, the multiplicative intercept parameter *ρ*, which primarily dictates the height of the development rate curve, can be modelled as a quadratic function of development stage. This structure was implemented by parameterizing a quadratic function of development stage, represented by consecutive integers, with the longest stage L4 represented by 0 (L2 = -2, L3 = -1, L4 = 0, L5 = 1, L6 = 2). Three points were used to derive an equation for the quadratic structure: (−2, *φy*_0_), (0, *y*_0_), and (2, *ψy*_0_). These points represent the values of *ρ* for the L2, L4 and L6 stages respectively. The *y*_0_ parameter represents the value of *ρ* at the longest development stage, L4. The *ψ* and *φ* parameters represent the ratio of *ρ*_*L*2_ and *ρ*_*L*6_, respectively, to *ρ*_*L*4_. This parameterization ensures that the values of *ρ* remain positive for all development stages. For a given integer representation of development stage *s*, the value of *ρ* is

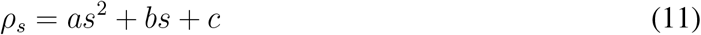

Plugging the above points into this equation and solving for the values of *a* and *b* gives

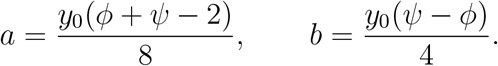

This strict quadratic relationship was relaxed by the addition of another set of multiplicative non-parametric adjustment parameters, 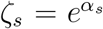 for *s* = 1, …, 5, to correct for the inflexibility of the quadratic curve in representing the relationship between *ρ* and development stage. The new value of the multiplicative intercept *ρ* at each stage of the model is 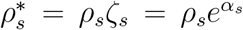 where the values of *α*_*s*_ have the distribution 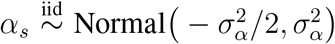 such that the expected value of 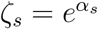 is 1.

### Bayesian Implementation

Effective methods for sampling from the posterior distribution of Bayesian models have been an active area in computational statistics for decades. Hamiltonian (or hybrid) Monte Carlo (HMC), proposed nearly 30 years ago (Neal, 1993), has recently achieved much greater popularity with the availability of convenient and powerful implementations. While Stan (Stan Development Team, 2020) is probably the most popular such tool, our model was written in Template Model Builder (Kristensen et al., 2015), which is primarily designed for maximum likelihood estimation. The R package tmbstan (Monnahan and Kristensen, 2018) was used to apply Stan’s No-U-Turn Sampler (NUTS), a particular step-size rule for HMC, to the model.

With the exception of the stage-structured component of the model, each parameter was sampled independently for each development stage. The parameters *H*_*A*_, *H*_*H*_ and *−H*_*L*_ were assigned Gamma priors, while the parameters *T*_*L*_ and *T*_*H*_ were assigned Normal priors. Although the values of *T*_*L*_ and *T*_*H*_ are strictly positive, they are large in magnitude so a Normal distribution with relatively small variance is sufficient. The scale parameters *σ*_*ϵ*_, *σ*_*υ*_ and *σ*_*α*_ were all assigned log-Normal priors. The *y*_0_ parameter of the quadratic curve effectively represents a development rate, so it was assigned a Gamma prior, while *φ* and *ψ* were both assigned Beta priors.

To assess the coverage and accuracy of the model, we performed a 10-fold cross-validation on both the semi-parametric model, a strictly parametric version of the model, and the MLE of the parametric model. The details of this procedure are outlined in the Supplementary Materials.

### Simulations

To obtain credible intervals for values that are of immediate use for decision making, we simulated insect development from posterior samples based on a year of hourly weather data from Timmins, Ontario where the wild colony was initially sampled. This data was obtained from the weathercan package in R (LaZerte and Albers, 2018). Since the scope of the model did not include overwintering larvae, each individual was assumed to begin the L2 stage at the same time, on April 1st, 2020. One thousand samples were taken from the posterior distribution, and one thousand individuals were generated from each of these samples. For each sample, the values of the parametric curves corresponding to each treatment temperature were calculated; to generate individuals, log-Normal *δ*_*i*_ values were sampled for each individual using the fitted scale parameter, *σ*_*ϵ*_. These *δ*_*i*_ values were multiplied by the population median response curve to obtain each individual response curve. Development was summed over each time step of the weather data based on each individual’s estimated development rate curve.

## Results

For the full, semi-parametric model, the MCMC sampler converged satisfactorily to the posterior distribution, as determined by a maximum 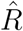 value of 1.003 from the rank-normalized samples (Vehtari et al., 2021). The number of samples was sufficient, with both tail and bulk effective sample sizes greater than 3200 (Vehtari et al., 2021), and there were no divergent transitions reported from the final samples (Stan Development Team, 2020). Some divergent transitions did occur in earlier stages of the model development, but we eliminated them by (1) reducing the sampler’s step size, (2) implementing Cauchy-distributed non-parametric adjustment parameters, and (3) decentralizing the model parameters in the model definition in order to make the posterior surface more amenable to HMC sampling (Gelman et al., 2020; Stan Development Team, 2020); decentralized model parameters are sampled from a standardized distribution and then shifted and scaled appropriately, rather than being sampled directly (Stan Development Team, 2020). The strictly parametric version of the model also converged satisfactorily with a sufficient effective sample size, returning no divergent transitions.

Prior predictive checks ensured that the resulting curves were reasonable; the prior distributions for parameter values that allowed unrealistic development rate curves were narrowed. Widening the priors for some parameters caused the sampler to report divergent transitions. The scale parameters *σ*_*ϵ*_ and *σ*_*υ*_ for the non-parametric adjustment components were particularly problematic, especially when their priors were concentrated on small values. Shifting the prior distributions upward eliminated these issues. The *σ*_*υ*_ parameter was poorly informed by the data; the posterior distribution of this parameter was close to its prior distribution. Widening the prior resulted in the same behaviour, and caused divergent transitions when the priors allowed for very small values of *σ*_*υ*_. Figure 2 demonstrates the behaviour of *σ*_*υ*_, along with the quadratic structure of the *ρ* parameter. It also shows the posterior distribution of another parameter of interest: the scale parameter for the distribution of individual variation, *σ*_*ϵ*_. This parameter also appears to have a quadratic relationship with developmental stage but in contrast to the *ρ* parameter, *σ*_*ϵ*_ is at its smallest in the L4 stage. This behaviour indicates that individuals’ development relative to the population median varies more in the early and late stages than in the middle stages of development. The upper right panel in Figure 2 shows the distribution of the multipliers at each stage for the non-parametric adjustment between the data and the parameterized quadratic structure across stages, as captured by the non-parametric adjustment term. These distributions are concentrated around one for all stages but L3, indicating that the data mostly support the quadratic structure. The model was also fitted without the quadratic structure; that is, each stage was fitted independently. The resulting posterior distributions were very similar to those from the stage-structured model formulation, indicating that the data support the quadratic parameterization.

**Figure 2:**
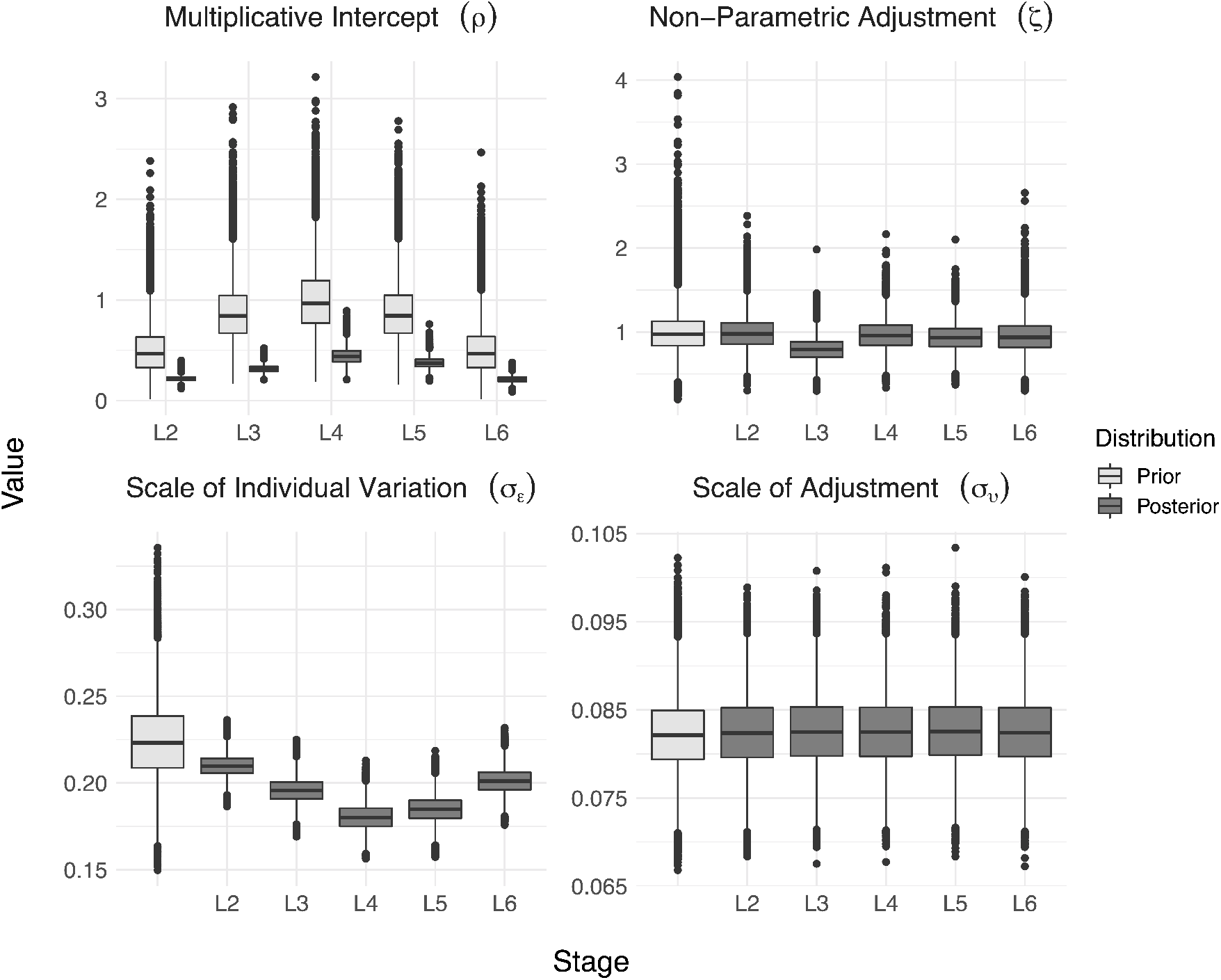
Box plots of prior and posterior distributions of *ρ, ζ, σ*_*ϵ*_ and *σ*_*υ*_ across developmental stages.

Figure 3 shows the mean absolute error between the log observations and the median log predictions for each observation, calculated from the cross-validation results. Since the observations are in days, which we would expect to have a skewed distribution, and since days to moult are on different scales for different temperatures, we are more interested in relative error than absolute error. For this reason, we calculate error on the log scale; we take the mean of the absolute log of the ratio between observation and prediction. A mean absolute log-scale error of zero (observed = predicted) is the best case, and larger error indicates poorer fit. As demonstrated by Figure 3, the fully parametric model consistently outperforms the semi-parametric model; the average log ratio rarely exceeds 0.1, which translates roughly to a 10% relative error. The unsustainable temperatures, both low and high, pose the most consistent problems for accurate estimation. Both models underperform for the 30 and 35^◦^C treatments in the L6 stage, which indicates some misfit of the parametric form to the data. Due to the unrestricted parameter spaces in the MLE method, many curves that were sampled contained negative development rates; these values were forced to zero, since negative rates are nonsensical within this framework. This pushed the upper boundaries of the prediction intervals for many observations to infinity, inflating the coverage probabilities and resulting in infinite log-scaled error values.

**Figure 3:**
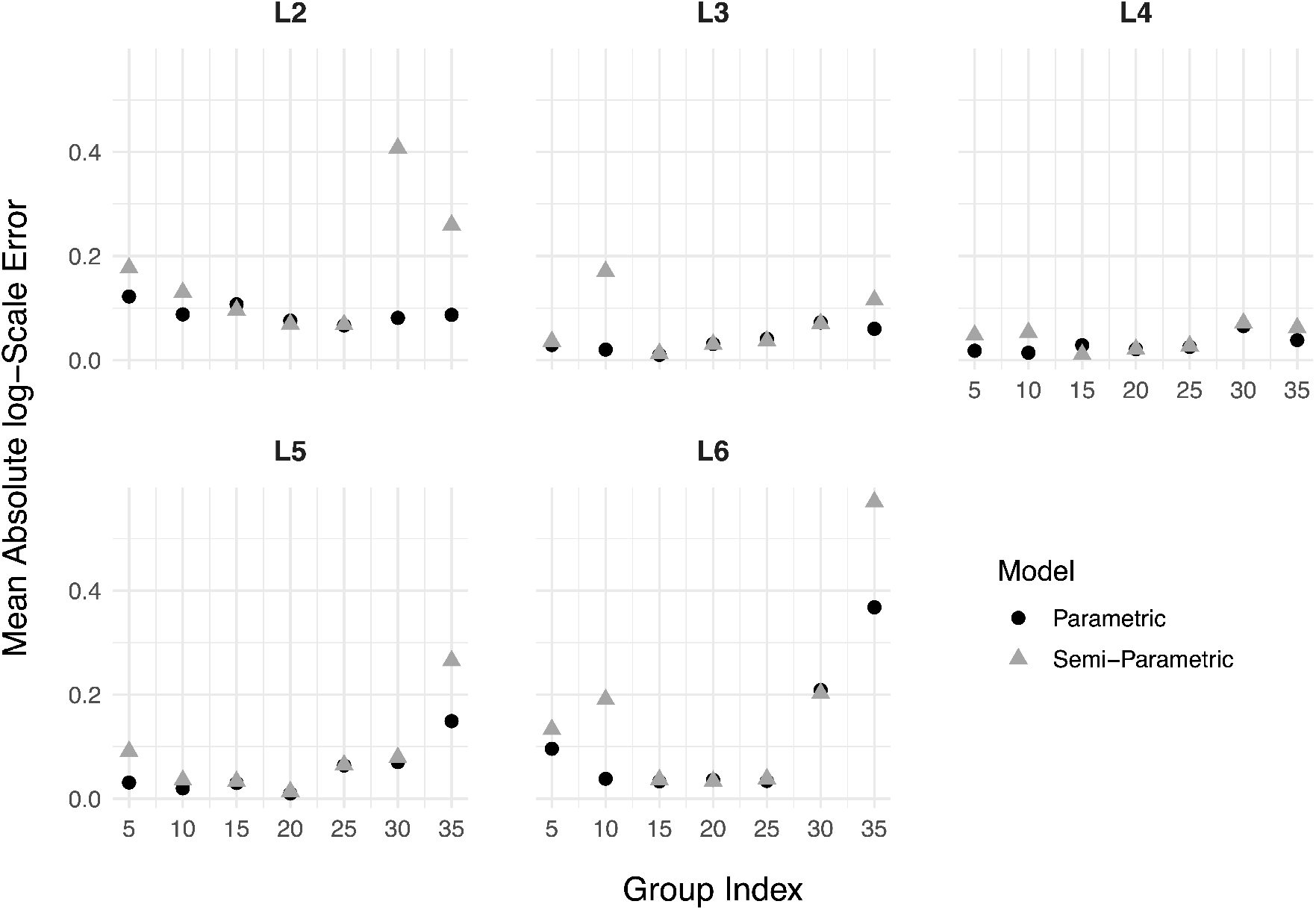
A comparison of the error of the semi-parametric and strictly parametric models across stages and development temperatures. The error equation is provided in the Supplementary Materials. Corresponding values for MLE method were infinite, and could not be included in the plot.

Figure 4 compares prediction intervals generated from the posterior samples of the strictly parametric model to the interval censored observations of development rate. The prediction intervals were created by sampling populations of individuals from the full posterior distribution; for each posterior draw, 200 values of the individual variation multiplier, *δ* were randomly sampled and multiplied by the development curve corresponding to their respective posterior draws. The range of resulting curves represents the full distribution of predictions. The central black curves represent the median development rates across all of the sampled individuals, while darkest ribbons represent the 50% prediction intervals, the next lightest represent the 95% prediction intervals and the lightest ribbons represent the 99% prediction intervals. The overlaid line segments in the plots represent the observed intervals of development rate. Since some of the lower bounds of the intervals of development time are zero, the upper intervals of the rates, the reciprocal of development time, go to infinity; these intervals are indicated by the small arrows at the top of the plots for the L3 and L4 stages. At the extreme temperatures for which transfer treatments were performed, the intervals are imputed with the posterior distribution of development rates at 20^◦^C using a moment matching technique (see Appendix). For the most part, the prediction intervals match the data well; in some instances, such as the upper temperatures in L2 and the lower temperatures in L3, there is some discrepancy between the model and the observed data. To investigate the source of the discrepancy, we fitted the model to a population generated from a central set of parameters from the prior distribution. There was no discrepancy between the prediction intervals and the “observed” data in this case, so the original discrepancy can likely be attributed to model misspecification, i.e. systematic differences between the data and the structure of our model.

**Figure 4:**
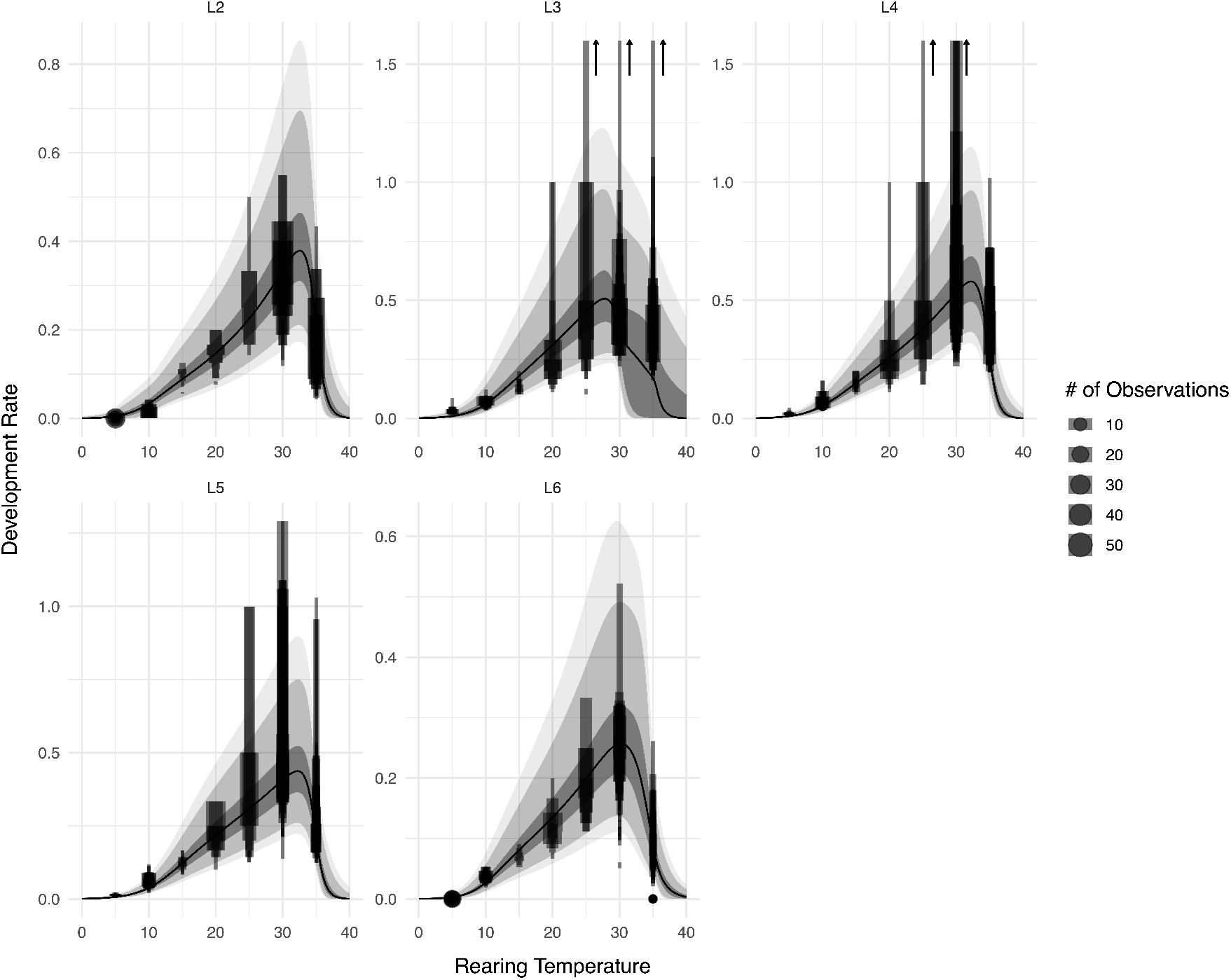
Ribbons represent prediction intervals generated from the posterior distribution; from darkest to lightest, they are 50%, 95% and 99% intervals. Black lines represent median predicted development rates. Black line segments represent observed development rates; rates are imputed for individuals reared at unsustainable temperatures using the method described in the Appendix.

The bottom panel of Figure 5 shows quantiles of daily development for the full simulated cohort. The vertical extent of the ribbons represents the credible intervals for the cohort’s developmental age over time, while the ribbons’ horizontal extents show the credible intervals of dates at which developmental milestones are reached. Though it uses a crude estimate of the cohort’s starting date for the L2 stage, the development timeline shown in the figure matches what we would expect from an Ontario budworm population in a typical year (Régnière et al., 2012). The black line in the upper panel of the figure shows mean daily temperature, while the background shading represents the expected development rate given temperature and proportion of individuals in each stage at a given date. This expected development rate is calculated using the median estimated development rate curve for each stage, weighted by the median estimated proportion of individuals in each stage on a given date. When the mean daily temperature line traverses darkly shaded areas (e.g. early June), budworm larvae are expected to develop rapidly.

**Figure 5:**
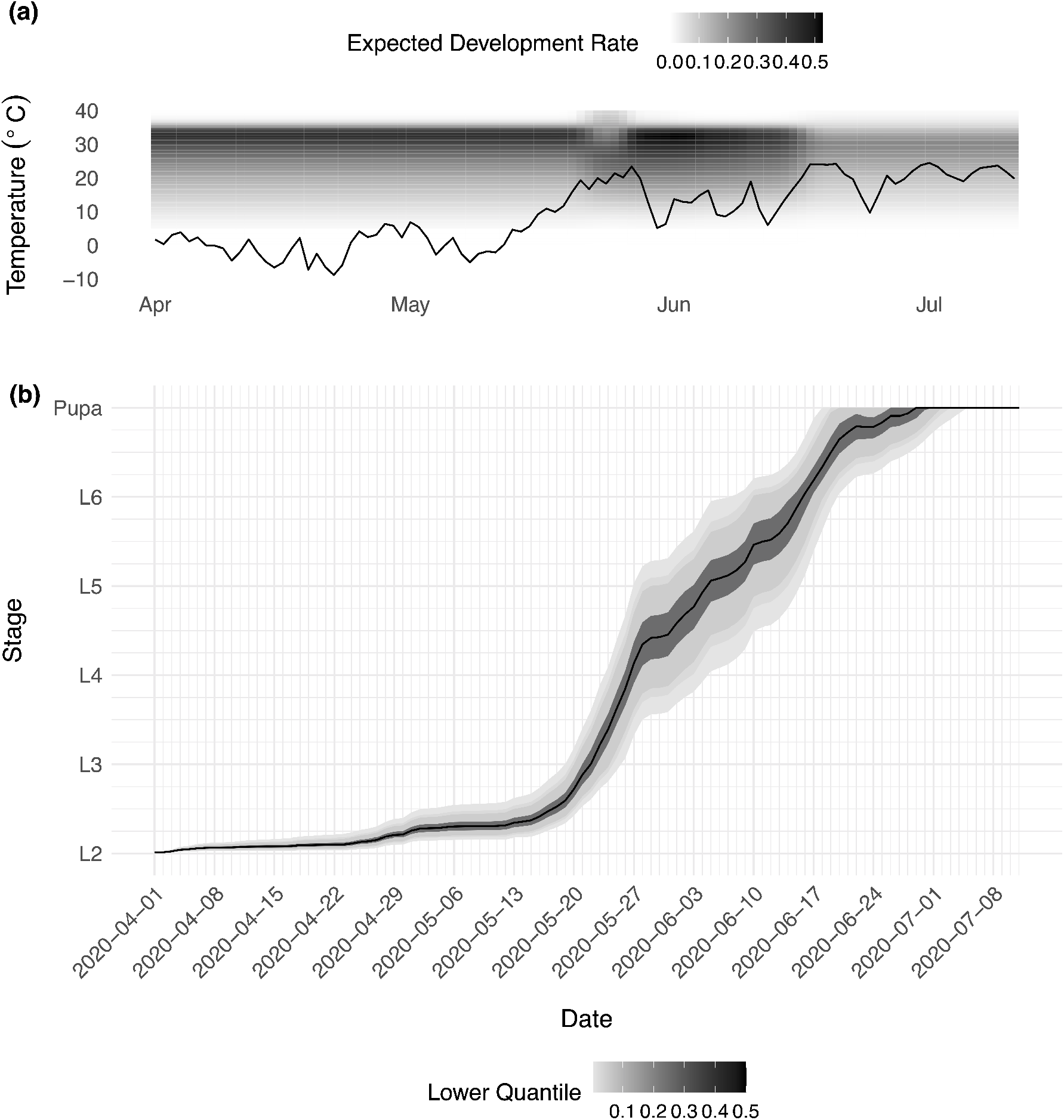
Top panel shows mean daily temperature in Timmins, ON for the year 2020. Background shading indicates expected development rate as a function of temperature and median proportions of the population in each development stage. Bottom panel shows symmetric credible intervals of development for the simulated population. The colours of the ribbons represent the lower percentile of the credible intervals.

## Discussion

### Posterior Samples

Our first objective was to obtain posterior samples from the model that could eventually be used for prediction. The Bayesian hierarchical model passed a range of standard tests indicating that the sampler had converged to the posterior distribution; the prior predictive checks showed that the prior distributions covered the desired span of development rate curves; and the posterior samples fit the data well, all suggesting that our sampled posterior distributions can be trusted for inference and prediction. We did need to modify our initial model specification to eliminate divergent transitions in the HMC sampler; model reparameterization, and adjustments of the prior distributions, are often necessary to obtain adequate sampling. (Such adjustments should only be done to address sampling issues, not to eliminate unwanted results!) Most of the sampling problems stemmed from overly restrictive prior specifications. For instance, when the prior on the scale parameter *σ*_*ϵ*_ was concentrated around values that were too small, the specified prior distribution of log(*δ*) = *E* was too narrow. Shifting the distributions of the scale parameters to larger values eliminated the issues with the sampler by allowing it to explore a wider range of *E* values. In general, overly large values of the scale hyperparameters (e.g. because of inappropriate priors) should be less problematic than overly small values; increasing the scale makes the likelihood surface of the higher-level parameter less steep. While in principle HMC can handle steep gradients in the posterior probability density, it generally fails when there is heterogeneity in steepness. The combination of small values of the scale parameters and wide-ranging prior distributions can cause the otherwise homogeneous posterior density to spike dramatically in certain regions.

For the non-parametric adjustment parameters *υ*_*j*_, large deviations from the estimated parametric curves in some ranges (e.g. development rates near the unsustainable temperatures) required thicker tails than provided by a Normal distribution. Using a Normal distribution for *υ*_*j*_, the posterior density of the curve parameters was highly bimodal, indicating that the model was sampling two distinct families of curves that agreed with the observations. Graphical exploration showed that the fits were being influenced by different sets of observations; drawing *υ*_*j*_ from a Normal distribution failed to allow enough flexibility around the parametric curve to match observed development rates across all of the experimental temperatures. Drawing *υ*_*j*_ from a Cauchy distribution instead resolved the problem.

### Stage Structure

Structure across development stages was incorporated into the model for the purpose of linking stages together, parameterizing the transformation of development rate curves across stages and creating a framework that could eventually permit a more sophisticated model, including relationships between other parameters across stages and correlation structure within individuals across stages. We assume that the baseline development rate (*ρ*) follows a quadratic curve across development stages (Régnière et al., 2012). The model also incorporates a non-parametric adjustment component (in the form of a *ζ*_*s*_ term) that permits departures from this structure; deviations of *ζ*_*s*_ from 1.0 indicate deviations of the development rate pattern from the quadratic skeleton. The posterior distributions of the deviation terms are concentrated around the null value of 1.0, confirming the assumption of a quadratic change in development rate across stages. The *ζ*_*s*_ parameter describing the L3 stage is centred at a value slightly less than 1.0, indicating that the estimated value of the *ρ* parameter at this stage is slightly larger than would be expected from a true quadratic relationship.

Overall, the results indicate that the posterior estimates of the curve of *ρ* across developmental stages are close to quadratic. This was reflected in the results of the model with stages treated independently; the posterior distributions resulting from the two sets of model runs (structured and independent) were very similar. This similarity indicates that the predictions resulting from the two sets of runs would be similar, implying that including the stage structure in the model does not give much of a predictive advantage. Previous individual-based development rate models have fitted development stages independently, with no link between curve features across stages. However, development is a continuous process; development rate curves are more likely to change gradually as individuals progress through the stages. The structure in this model could eventually make way for a development surface rather than a development curve, with the added dimension coming from an individual’s current developmental state. Incorporating continuous proxies of development such as size or weight could further solidify the relationship between development and current stage. Although our discrete representation of development stage is an oversimplification, it is a useful placeholder until a continuous marker of development can be measured, observed and integrated into the model.

Although our model improves on current practice by allowing for the pattern of development rates across stages at the population level, it still assumes independence in the development rates of individual larvae across stages. Little is known about the validity of this assumption, which is ubiquitous among individual-based development rate models. One can easily imagine that across-stage correlations could be positive (due e.g. to genetic variation among individuals) or negative (due to compensatory effects, where slow-growing individuals are able to catch up in later stages). The current model could be expanded to allow such within-individual correlations, relaxing the assumption of independence across stages and providing a better assessment of variability at the population level.

### Non-Parametric Adjustment

Including non-parametric adjustment parameters for each of the parametric curves in the model (the development rate curve, Eq. 10, and the quadratic curve of development rate by stage, Eq. 11) relaxes the assumption that these phenomena conform strictly to a particular mathematical relationship. In this framework, the parametric model acts as a baseline or “skeleton” for the observed variation in development rate across temperature or stages; the non-parametric adjustment parameters add flexibility, allowing the model to accommodate departures from the parametric skeleton. The non-parametric adjustment framework has two advantages over the alternative fully non-parametric approach (e.g. using splines or Gaussian process models to model the TPC or development curve across stages). Unlike fully non-parametric models, the parametric skeleton provides interpretable parameters that can be connected back to underlying physiological or ecological processes. While these parameters are less useful when the parametric model is a poor fit to the data, they are still the best available estimates, and provide information that is lost in a fully non-parametric model. The parametric skeleton also facilitates the specification of prior distributions. While one could in principle generate informed prior distributions for non-parametric curves (e.g. by generating a sample of curves from a non-parametric family and having subject-area experts rank them for plausibility), it is much easier to base priors on previously estimated parameters from mechanistic models of insect development. In theory, this component of the model should reduce the bias introduced by imposing a strict parametric form on the insects’ development. However, the cross-validation study showed that the predictions were in fact more accurate without this model component; this result can likely be attributed to overfitting. The additional flexibility in the model causes it to conform more closely to the data, but with small or noisy data sets this flexibility can also result in poor generalization, i.e. models that do not fit the validation data well. In our case, the parametric model is more robust to small sample sizes and is therefore preferable for prediction. For both model types, the Bayesian fitting method outperformed the MLE, due to the naive assumption of the parameters’ multivariate normality. This assumption led to unrealistic sampled parameter values, resulting in infinite prediction error. The regularization of estimates provided by the Bayesian method solves this issue, preventing the model from providing unrealistic or nonsensical estimates.

### Simulations

We can obtain credible intervals for values of interest by sampling populations and individuals directly from the posterior distribution and using these values for simulations based on realistic temperature profiles (Figure 5). Current simulations using weather data to predict insect population development rely on point estimates of model parameters, with the only stochastic component coming from individual-level variation around a deterministic development curve (Régnière et al., 2014). This simulation method completely neglects uncertainty around parameter estimates, and therefore underestimates the uncertainty in downstream predictions. In contrast, using Bayesian methods to sample both individual- and population-level variation from the estimated posterior distributions propagates the full range of model uncertainty through the simulations, so that they are reflected in the credible intervals for predicted values. This technique is equally applicable to scenarios informed by current conditions; based on historical weather data; or using climate projections to predict how insect populations will develop in the future. Predictions and projections from a fitted model cannot be more reliable than the data they are based upon. The data used to fit this model was collected in laboratory rearing conditions, from insects kept under constant temperature regimes. In nature, insects feed on a natural diet and are subject to highly variable (and heterogeneous) temperature conditions. Going forward, our predictions should be validated against insect development observed in the wild. Future empirical work includes experiments that use more realistic diets and temperature regimes in the laboratory, enabling us to use our laboratory data to predict insect phenology in the wild.

## Conclusions

Our Bayesian hierarchical model provided more consistent fits to the data than were achieved by MLE; CIs on the development curve estimates based on the MLE often included biologically impossible values. Because the full model that incorporated non-parametric adjustment components was prone to overfitting, the parametric version of the model was used for real-weather simulations. The Bayesian prediction intervals on the development curve estimates excluded implausible values and fit the data well. This demonstrates the advantage of parameter regularization, which is achieved using Bayesian methodology. The incorporation of a functional representation of the transformation of development rate curves across stages lays the groundwork for future models to treat this transformation as a more continuous process, and to incorporate the correlation of individual development across stages. Using the full posterior distribution to generate weather-driven simulations of population development propagates the uncertainty around parameter estimates to downstream estimates of a population’s developmental timing. These simulations ultimately provide more realistic and robust estimates than those generated from single point estimates.

## References

Balakrishnan, N. and Leung, M. (1988). Order statistics from the type i generalized logistic distribution. Communications in Statistics-Simulation and Computation, 17(1):25–50.

Bean, J. (1961). Predicting emergence of second-instar spruce budworm larvae from hibernation under field conditions in Minnesota. Annals of the Entomological Society of America, 54(2):175–177.

Bewick, S., Cantrell, R. S., Cosner, C., and Fagan, W. F. (2016). How resource phenology affects consumer population dynamics. The American Naturalist, 187(2):151–166.

Blais, J. R. (1983). Trends in the frequency, extent, and severity of spruce budworm outbreaks in eastern Canada. Canadian Journal of Forest Research, 13(4):539–547.

Bolker, B. M. (2008). Ecological models and data in R. Princeton University Press.

Calle, M. L. (2002). The Analysis of interval censoring and double censoring via Markov chain Monte Carlo methods. Technical report, Universitat de Vic - Central University of Catalonia.

Cameron, D., McDougall, G., and Bennett, C. (1968). Relation of spruce budworm development and balsam fir shoot growth to heat units. Journal of Economic Entomology, 61(3):857–858.

Candau, J.-N., Dedes, J., Lovelace, A., MacQuarrie, C., Perrault, K., Roe, A., Studens, K., and Wardlaw, A. (2019). Validation of a spruce budworm phenology model across environmental and genetic gradients: applications for budworm control and climate change predictions. Technical report, SERG International.

Ceglar, A., Črepinšek, Z., Kajfež-Bogataj, L., and Pogačar, T. (2011). The simulation of phenological development in dynamic crop model: The Bayesian comparison of different methods. Agricultural and Forest Meteorology, 151(1):101–115.

Chuine, I. and Régnière, J. (2017). Process-based models of phenology for plants and animals. Annual Review of Ecology, Evolution, and Systematics, 48:159–182.

Crimmins, T. M., Gerst, K. L., Huerta, D. G., Marsh, R. L., Posthumus, E. E., Rosemartin, A. H., Switzer, J., Weltzin, J. F., Coop, L., Dietschler, N., et al. (2020). Short-term forecasts of insect phenology inform pest management. Annals of the Entomological Society of America, 113(2):139–148.

Damos, P. and Savopoulou-Soultani, M. (2012). Temperature-driven models for insect development and vital thermal requirements. Psyche, 2012. Article ID 123405.

Dennis, B., Kemp, W., and Beckwith, R. (1986). Stochastic model of insect phenology: estimation and testing. Environmental Entomology, 15(3):540–546.

Fu, Y. H., Campioli, M., Van Oijen, M., Deckmyn, G., and Janssens, I. A. (2012). Bayesian comparison of six different temperature-based budburst models for four temperate tree species. Ecological Modelling, 230:92–100.

Gelman, A., Vehtari, A., Simpson, D., Margossian, C. C., Carpenter, B., Yao, Y., Kennedy, L., Gabry, J., Bürkner, P.-C., and Modrák, M. (2020). Bayesian workflow. arXiv preprint 2011.01808.

Hudes, E. and Shoemaker, C. (1988). Inferential method for modeling insect phenology and its application to the spruce budworm (Lepidoptera: Tortricidae). Environmental Entomology, 17(1):97–108.

Ikemoto, T. (2005). Intrinsic optimum temperature for development of insects and mites. Environmental Entomology, 34(6):1377–1387.

Kristensen, K., Nielsen, A., Berg, C. W., Skaug, H., and Bell, B. (2015). TMB: automatic differentiation and Laplace approximation. arXiv preprint 1509.00660.

Lawrence, R. K., Mattson, W. J., and Haack, R. A. (1997). White spruce and the spruce budworm: defining the phenological window of susceptibility. The Canadian Entomologist, 129(2):291– 318.

LaZerte, S. E. and Albers, S. (2018). weathercan: Download and format weather data from Environment and Climate Change Canada. The Journal of Open Source Software, 3(22):571.

Lemoine, N. P., Burkepile, D. E., and Parker, J. D. (2014). Variable effects of temperature on insect herbivory. PeerJ, 2:e376.

Lysyk, T. (1989). Stochastic model of eastern spruce budworm (Lepidoptera: Tortricidae) phenology on white spruce and balsam fir. Journal of Economic Entomology, 82(4):1161–1168.

McCarthy, M. A. and Masters, P. (2005). Profiting from prior information in bayesian analyses of ecological data. Journal of Applied Ecology, pages 1012–1019.

McManis, A. E., Powell, J. A., and Bentz, B. J. (2018). Developmental parameters of a southern mountain pine beetle (Coleoptera: Curculionidae) population reveal potential source of latitudinal differences in generation time. The Canadian Entomologist, page 1.

Monnahan, C. and Kristensen, K. (2018). No-U-turn sampling for fast Bayesian inference in ADMB and TMB: Introducing the adnuts and tmbstan R packages. PloS ONE, 13(5).

Neal, R. M. (1993). Probabilistic inference using Markov chain Monte Carlo methods. Department of Computer Science, University of Toronto Toronto, ON, Canada.

Perrault, K., Wardlaw, A., Candau, J.-N., Irwin, C., Demidovich, M., MacQuarrie, C., and Roe, A. (2021). From branch to bench: establishing wild spruce budworm populations into laboratory colonies for the exploration of local adaptation and plasticity. The Canadian Entomologist, 153(3):374–390.

Porter, J., Parry, M., and Carter, T. (1991). The potential effects of climatic change on agricultural insect pests. Agricultural and Forest Meteorology, 57(1-3):221–240.

Pruess, K. P. (1983). Day-degree methods for pest management. Environmental Entomology, 12(3):613–619.

Qiu, T., Song, C., Clark, J. S., Seyednasrollah, B., Rathnayaka, N., and Li, J. (2020). Understanding the continuous phenological development at daily time step with a Bayesian hierarchical space-time model: impacts of climate change and extreme weather events. Remote Sensing of Environment, 247:111956.

Quinn, B. K. (2017). A critical review of the use and performance of different function types for modeling temperature-dependent development of arthropod larvae. Journal of Thermal Biology, 63:65–77.

Rebaudo, F. and Rabhi, V.-B. (2018). Modeling temperature-dependent development rate and phenology in insects: review of major developments, challenges, and future directions. Entomologia Experimentalis et Applicata, 166(8):607–617.

Régnière, J. (1984). A method of describing and using variability in development rates for the simulation of insect phenology. The Canadian Entomologist, 116(10):1367–1376.

Régnière, J., St-Amant, R., and Duval, P. (2012). Predicting insect distributions under climate change from physiological responses: spruce budworm as an example. Biological Invasions, 14(8):1571–1586.

Régnière, J. (1987). Temperature-dependent development of eggs and larvae of Choristoneura fumiferana (Clem.) (Lepidoptera: Tortricidae) and simulation of its seasonal history. The Canadian Entomologist, 119(7-8):717–728.

Régnière, J., Powell, J., Bentz, B., and Nealis, V. (2012). Effects of temperature on development, survival and reproduction of insects: experimental design, data analysis and modeling. Journal of Insect Physiology, 58(5):634–647.

Régnière, J., Saint-Amant, R., Béchard, A., and Moutaoufik, A. (2014). BioSIM 10: User’s manual. Laurentian Forestry Centre.

Schoolfield, R., Sharpe, P., and Magnuson, C. (1981). Non-linear regression of biological temperature-dependent rate models based on absolute reaction-rate theory. Journal of Theoretical Biology, 88(4):719–731.

Stan Development Team (2020). Stan Modeling Language Users Guide and Reference Manual 2.25.

Stedinger, J., Shoemaker, C., and Tenga, R. (1985). A stochastic model of insect phenology for a population with spatially variable development rates. Biometrics, 41(3):691–701.

Sturtevant, B. R., Achtemeier, G. L., Charney, J. J., Anderson, D. P., Cooke, B. J., and Townsend, P. A. (2013). Long-distance dispersal of spruce budworm (Choristoneura fumiferana Clemens) in Minnesota (USA) and Ontario (Canada) via the atmospheric pathway. Agricultural and Forest Meteorology, 168:186–200.

Thorsen, S. and Höglind, M. (2010). Modelling cold hardening and dehardening in timothy. Sensitivity analysis and Bayesian model comparison. Agricultural and Forest Meteorology, 150(12):1529–1542.

Uvarov, B. (1931). Insects and climate. Transactions of the Royal Entomological Society of London, 79(pt. 1).

Vehtari, A., Gelman, A., Simpson, D., Carpenter, B., and Bürkner, P.-C. (2021). Ranknormalization, folding, and localization: An improved R for assessing convergence of MCMC. Bayesian Analysis, 16(2):667–718.

Wardlaw, A., Perrault, K., Roe, A., Dedes, J., Irwin, C., MacQuarrie, C., and Candau, J.-N. (2022). Methods for estimating and modelling spruce budworm development rates at constant temperatures. The Canadian Entomologist, 154.

