## Supplementary Materials for "Predicting the temperature-driven development of stage-structured insect populations with a Bayesian hierarchical model"

### Supplementary Materials: Development Rate Imputation

Due to the potential for high mortality outside of rearing temperatures in the "sustainable" range, individuals reared at these unsustainable temperatures were subject to transfer treatments. Since a proportion of each stage is spent at 20°C under this regime, it is impossible to observe development at the unsustainable temperatures directly. For the purpose of comparing observations to model fits, "observed" development rates at the unsustainable temperatures need to be imputed. If we re-write Equation 1 from the main text for an individual $i$ who has completed a development stage, we have

$$1 = \delta_i(r_l^* t_{l,i} + r_s^* t_{s,i}) \tag{1}$$

where $r_l^*$ and $r_s^*$ are the population median development rates at the lethal and sustainable temperatures, and $t_{l,i}$ and $t_{s,i}$ are time in days spent at those temperatures. The values of $t_{l,i}$ and $t_{s,i}$ are known, and $r_s^*$ can be estimated from the subpopulation that is reared at the corresponding sustainable temperature ($T_s$). The values $\delta_i$ and $r_l^*$ are both unknown, and our value of interest, the development rate of individual $i$ at the lethal temperature $T_l$ is equal to $\delta_i r_l^*$. Rearranging this equation gives

$$\delta_i = (r_l^* t_{l,i} + r_s^* t_{s,i})^{-1} \tag{2}$$

By design, $\delta_i$ is a log-Normally distributed random variable such that $\log(\delta_i)$ has mean 0; thus, the median value of $\delta_i$ is expected to be 1. We can therefore approximate $r_s^*$ by setting

$$1 = \text{median}(\delta) \approx \text{median}\big((r_l^* t_l + r_s^* t_s)^{-1}\big) \tag{3}$$

This equation can be solved for $r_s^*$ using the `uniroot` function in R. Upon solving for $r_s^*$, we can calculate the $\delta_i$ values for each individual, and multiply them by the estimate of $r_s^*$ to obtain the estimated development rates for each individual reared at lethal temperatures.

### Supplementary Materials: Cross-Validation

To assess the coverage and accuracy of our model, we performed a 10-fold cross-validation. Both the semi-parametric model and a strictly parametric version of the model were fitted to each fold (subsample) of the data. To ensure that each subpopulation was properly represented, the data was stratified by rearing temperature, and individuals were randomly assigned to cross-validation folds. We fitted the model to all individuals *not* included in each fold and compared the predicted values to the observations for the held-out individuals. We took 14,000 samples from the posterior distributions, and generated data sets of 1,000 individuals from each draw. To compare the model fits directly to the observed data, we simulated the sampled individuals' development under the treatment conditions of each unique observation. In the sustainable temperature treatments, the number of days to moult was calculated for each simulated individual based on their respective development parameters; in unsustainable temperature treatments, we held simulated individuals at the treatment temperature for the same amount of time as each unique observed insect, and then calculated the amount of time it would take each simulated individual to complete the development stage at 20°C. We calculated the 5th, 50th and 95th quantiles of these values for all the simulated individuals; the 5th and 95th quantiles represent a 90% prediction interval for each observation. A similar procedure was performed using maximum likelihood estimation. The parametric model was fitted to each withheld subset of data, and confidence intervals were obtained using the methods described in Bolker (2008). Maximum likelihood estimates were found using R's `optim` function on the TMB model code with 100 different starting values; from these 100 optimizations, the parameter set that minimized the negative log likelihood was taken as the MLE. We then sampled from a multivariate normal distribution parameterized by these fitted values, analogous to the posterior samples we obtained from the Bayesian methods; once these samples were taken, the remainder of the procedure was identical. To assess coverage, the observed intervals of days to moult were compared with their respective 90% prediction intervals. If the intervals overlapped, we concluded that the observation fell inside the prediction interval. We would expect this method to slightly overcover the observations, since there is a possibility that the true, uncensored number

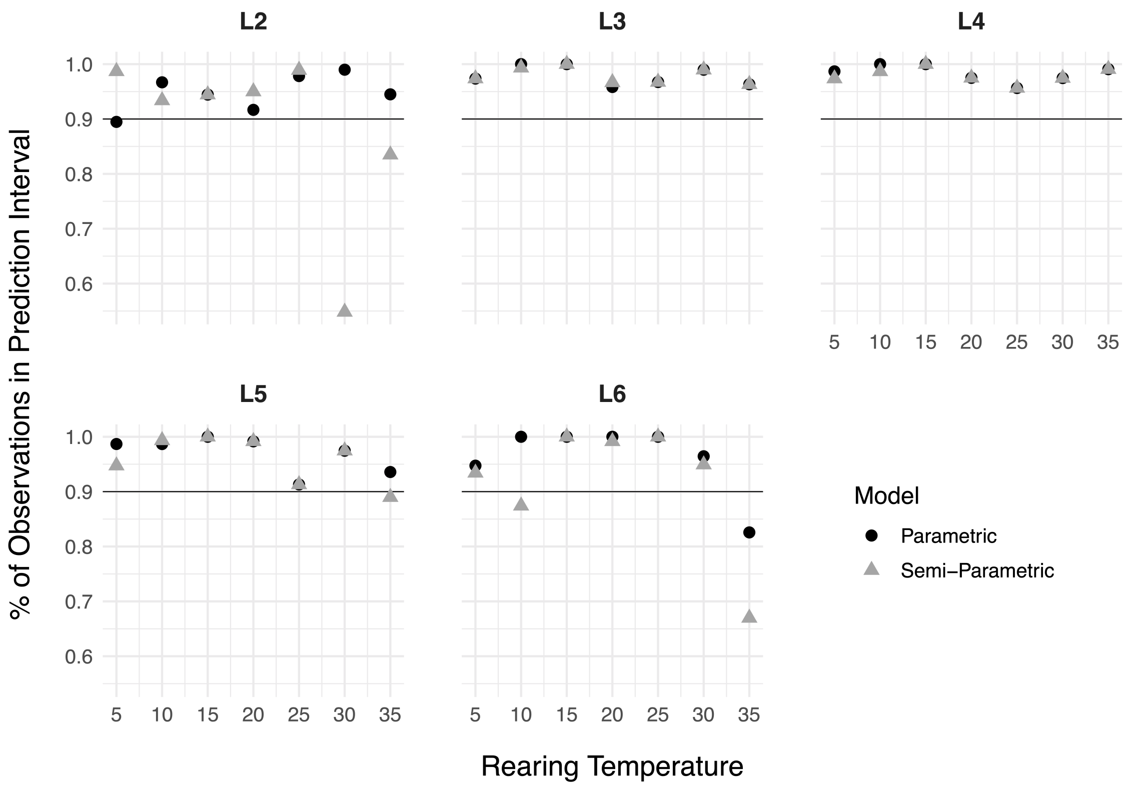


Coverage probabilities across stages and rearing temperatures for both Bayesian model types.

of days to moult could fall outside the prediction interval, even if the two intervals overlapped. The figure shows the coverage probabilities for each rearing temperature and development stage, and for each model type. To measure the accuracy of the model predictions, we compared each observation to the population median prediction. Since we are more interested in relative error of development times than absolute error, we measured error on the log scale. For each rearing temperature $T$, each stage $s$ and each model $m$, the error was calculated as

$$\text{MAE} = \frac{1}{N}\sum_{i=1}^{N} |\log(y_{Ts_i}) - \log(\hat{\mu}_{Tsm_i})| \tag{4}$$

where $y_{Ts_i}$ is the observed number of days to moult, and $\hat{\mu}_{Tsm_i}$ is the median prediction for each observation. To deal with the interval censored data, the observed value was reported as the interval's upper or lower bound, depending on whether the prediction was above or below the interval; if the prediction fell within the interval, the ratio was reported to be 1.

### Supplementary Materials: Posterior Distribution

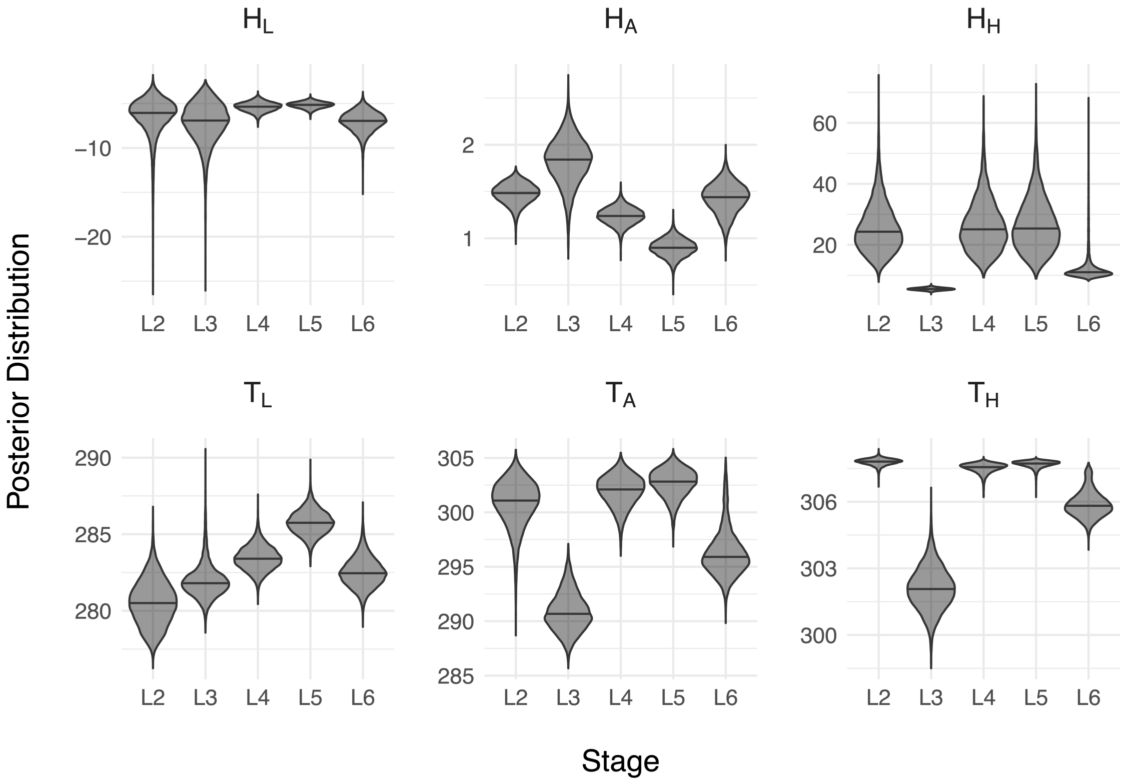


Posterior distributions of development curve parameters across developmental stages.
